# The role of symmetry breaking in connectivity and neurodegeneracy in the brain’s resting state dynamics

**DOI:** 10.1101/2025.02.10.637533

**Authors:** Kashyap Gudibanda, Jan Fousek, Spase Petkoski, Viktor Jirsa

**Affiliations:** Aix Marseille Univ, INSERM, INS, Inst Neurosci Syst, Marseille, France

## Abstract

The functioning of brain networks can be broadly categorized as an interplay of two main contributors, namely the neurological processes dictating the local dynamics and the patterns of anatomical connections that enable the interactions between the various local processes, resulting in a global emergent behavior. Using Brain Network Modeling we describe the constraints of the anatomical structure upon the resting state dynamics of the human brain. We identify a low-dimensional representation of the brain states, the Resting State Manifold (RSM), and leveraging network degeneracy and its relationship to structural properties, we identify dynamic patterns supported by the RSM. Noise driven dynamics of the BNM are used to explore the manifold and validate the contours of degeneracy. We demonstrate that the patterns of degeneracy regulate the dynamics of the broader system in the presence of external and/or internal perturbations, revealing the productive relationship between emergent network processes and their constituent network entities.

## Introduction

While there is an extraordinary diversity to the phenomena exhibited by the human brain, the dynamical repertoire associated with its resting state has garnered significant attention.^1–6^. Considerable effort has been invested in modeling the contribution of the anatomy to what is essentially a spatio-temporal phenomenon.^7–11^ These works have attempted to discern the underlying mechanisms of the resting state dynamics using a combination of dynamical systems theory, statistics and computational techniques, often through the framework of virtual brains^12,13^. Under the umbrella of resting state dynamics, a wide variety of attributes have been studied in the contexts of neurodegenerative disorders^14–16^, aging^17–19^, consciousness^20–22^, psychiatric disorders^23^, etc. However, the evolution of these dynamics, resulting from transitions between different brain states is key to having a complete description of the dynamical repertoire.

In this work, we adopt the brain network modeling approach, also known as virtual brains,^24,25^, where in absence of other hypothesis for cortical heterogeneity the network is constructed by setting up identical neural masses held together by a structural connectome that is assumed to be static at the time-scales on the neural activity occurs^9,26^. The activity of the BNM is generated by the interactions between the brain regions (neural masses), which receive specified inputs from their respective neighbours^12,13,27^. The simulated output of the BNM resides in a very high-dimensional space and is typically quantified using statistical (such as functional connectivity) and/or graph-theoretic metrics^8,28–30^. The BNM is then validated by comparing the metrics evaluated on the simulated activity to those calculated on recorded/measured activity^9,12,18,28,30–34^.

In our BNM framework, due to the dynamical repertoire of the chosen model for the mesoscopic neuronal activity^35^, the fixed points of the dynamical system are considered to be the brain states. Hence, spontaneous activity mandates the presence of either external currents or noise^7,9,13,32,36,37^for tracing out a trajectory in the space of possible brain states. We utilize this aspect to define a low-dimensional manifold on which the states are organized according to a simple scalar metric. To this end, we employ the Laplacian form of the structural connectivity matrix^38–40^. The Laplacian space is chosen primarily for the purpose of dimensionality reduction. However, it also has the added advantage that its eigenbasis extracts a hierarchy of spatial organizations from the connectome, thereby allowing us to identify the activation of specific subnetworks.

With a well-defined manifold, we tap into the notion of degeneracy to understand and predict the dynamics that can be supported by the manifold. In the neurobiological context, degeneracy is defined as the ability of non-identical structural elements to perform identical functions or result in identical outputs^41–45^. Specifically, we are interested in dynamical/network degeneracy, where an identical network function is performed by a broad set of network configurations with different parameters^46–49^. We use this framework of network degeneracy to quantify the effect of asymmetry introduced by the structural connectome. An interpretation of the degeneracy in terms of the robustness of network function allows us to characterize the structured flow on the low-dimensional resting state manifold. Structured Flows on Manifolds (SFMs) denote manifolds and their associated flows. A limit cycle, for instance, comprises a closed orbit with a flow driving the state through the manifold. More complex SFMs comprise higher-dimensional manifolds and multistable flows^50,51^. In that sense, we are building upon earlier works on the role of asymmetric couplings in the emergence of low-dimensional attractive manifolds and the flows thereupon^10,52^. These result in attractive self-organization functional subspaces that are relevant for the behavior^53^, and are related to self-organization in brain networks and moreover, information entropy and free energy^54^ within an information-theoretic framework^50^.

## Results

We construct a Brain Network Model (BNM) to study the role of symmetry-breaking due to structural connectivity. The local dynamics in the network model are simulated by a 2D Neural Mass Model (NMM), derived from an ensemble of all-to-all coupled QIF neurons, and the brain states correspond to the stable fixed points of the system, encompassing its dynamical repertoire, as also confirmed by our extensive numerical simulations. We build a low dimensional representation of these brain states from the Laplacian of the connectivity matrix and their distribution is mapped in the framework of parameter-degeneracy in the system. A noise-driven exploration of the brain states is employed to validate the contours of degeneracy and the consequential flow on the low dimensional manifold. We also provide a geometric interpretation of the degeneracy and a possible expansion to include symmetry-breaking by regional heterogeneity.

### Brain Network Model

The BNM is constructed as described in Methods-1. The local dynamics are modeled by the NMM^35^, which describes the evolutions of the firing rate (*r*) and membrane potential (*v*). This model has already been used in virtual brains to explain the emergence of various dynamical features relevant for neuroimaging at different time-scales^10,55^, as well as for explaining the compensation effects during aging^18^ and the change in brain’s dynamical working point during different stages of consciousness^22^. Moreover, the model was used to explore different strategies for tackling the non-identifiability in the pathology-linked degradation of brain’s connections^34^. The parameters of the single network node are chosen such that the system exhibits bistability, with a stable focus for a high firing rate and a stable node for a low firing rate. These are used to define our *up* and *down* states respectively. The network dynamics arises from interacting bistable nodes, where the oscillatory dynamics of the up-state provide the fast oscillations and set the characteristic frequency of the system. The slow oscillations in the network arise from noise-induced transitions between the up and down states (via the unstable fixed point) of the network nodes. A representative quantifier of the system behavior thus becomes the occupancy of the up-state of the network. In Fig. 1A, we show the nullclines (in red), the basin of attraction (in black), and the phase-flows (as arrows). Sample trajectories of the system, in the vicinity of both the stable focus and node, are also shown in blue.

**Figure 1.**
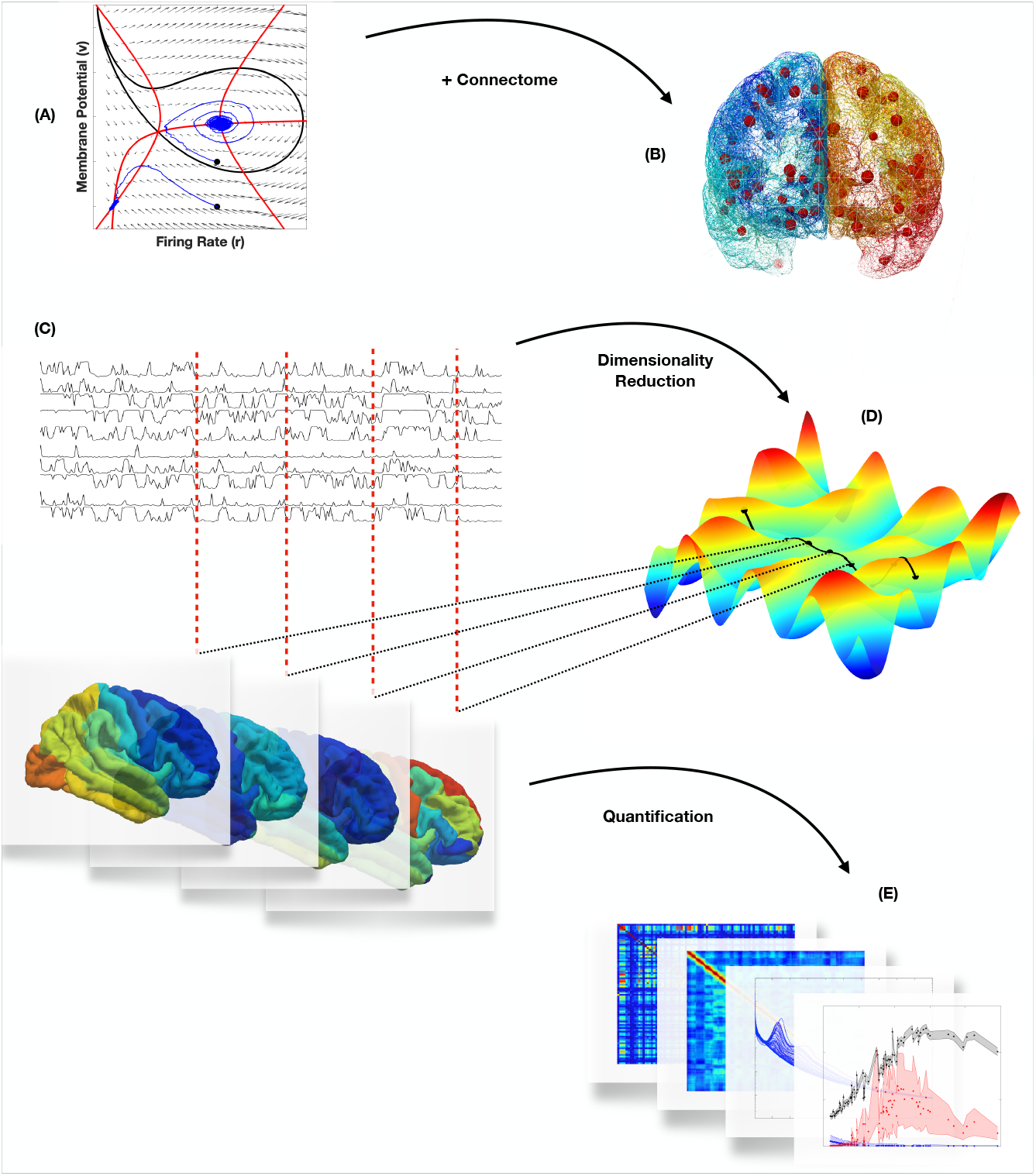
Schematic summary. (A) A non-linear system with bi-stability is selected to model the dynamics of the neural mass. In our case, the system has two stable fixed points, where the stable node is denoted as the down-state and the stable spiral as the up-state. (B) A brain network model, comprised of these interconnected neural masses, is constructed using a representative structural connectome from the HCP dataset. (C) A stochastic simulation of the network model gives rise to complex switching dynamics, as a result of symmetry-breaking induced by the connectome, and generates a high dimensional output. (D) The output of the BNM is transformed into a low-dimensional manifold on which the system is seen as tracing a trajectory while exploring different brain states. (E) Exploration of the manifold, seen as transitions between brain states, is studied in the framework of parameter degeneracy.

### Laplacian Eigenbasis

The Laplacian form of the structural connectivity matrix plays a key role in the definition of a low-dimensional representation of brain states. Decomposing the symmetric normalized form of the Laplacian matrix into its eigen-basis extracts the hierarchy of spatial organizations, that are manifested as structural partitions. Surface plots of the leading vectors are plotted and discussed in Fig. 2. We exploit this hierarchy to define a low-dimensional embedding and extract the resting state manifold. The symmetric normalized form also ensures that the eigenvalues are real, non-negative and bound in the interval [0, 2], expanding the comparability of results across various connectomes.

**Figure 2.**
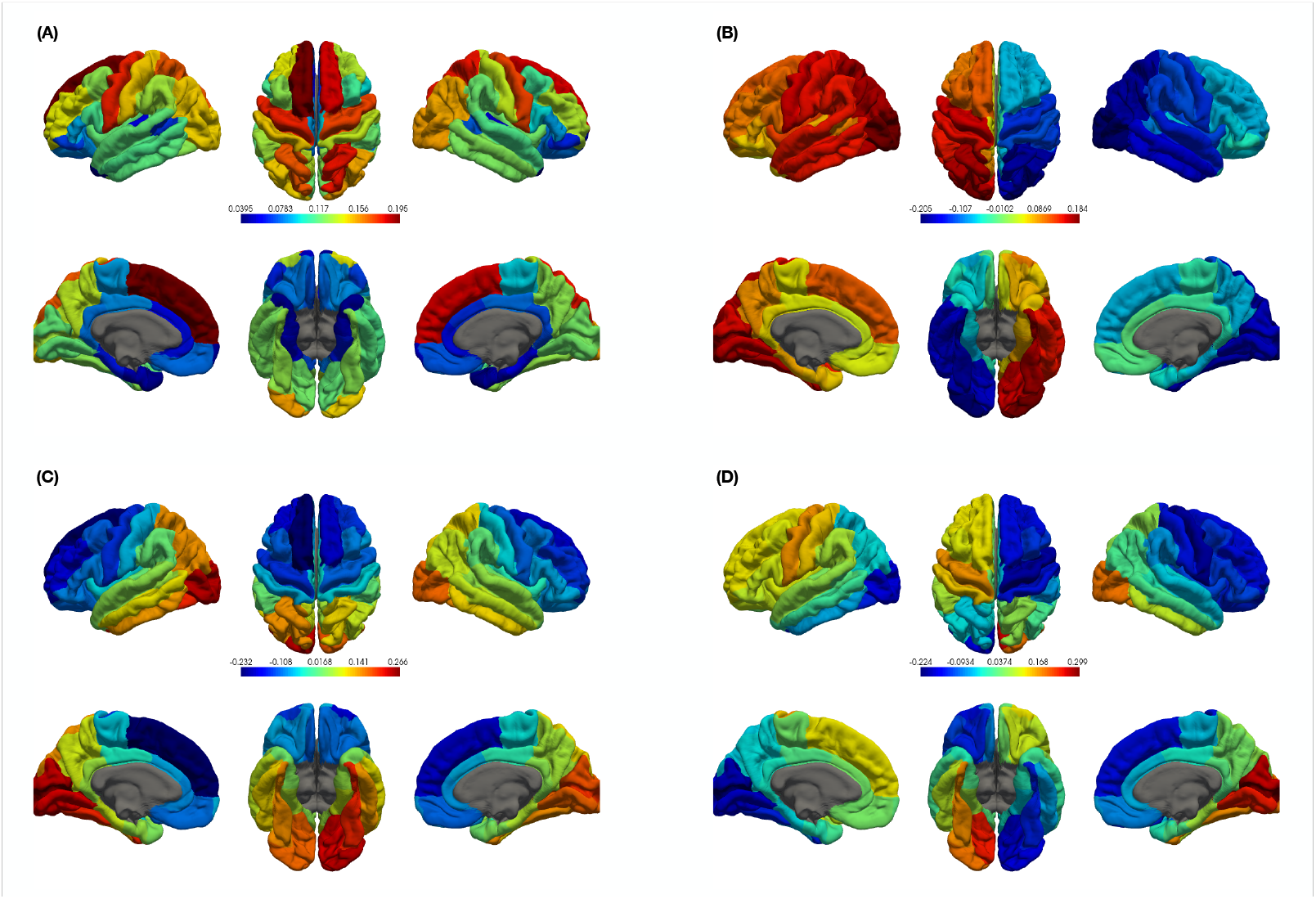
Eigenvectors of the Laplacian matrix. The four leading eigenvectors of the Laplacian matrix (*L*) are shown as surface plots. (A) The leading eigenvector (*e*_0_) corresponding to the zero eigenvalue. The coefficients of (*e*_0_) are proportional to the in-strengths of the regions. Since the network is connected, *λ*_0_ = 0 is unique and the coefficients of *e*_0_ are all of the same sign, indicating that the largest structural organization in the network is the network itself. (B) The second eigenvector (*e*_1_), corresponding to the first non-zero eigenvalue *λ*_1_ = 0.1016. *e*_1_ extracts the largest partition in the connectome, that is the separation of the hemispheres. The left and right hemispheres are separated by positive and negative coefficients of *e*_1_ respectively. We ignore *e*_2_ as it only highlights the cerebellum and gives no further information. (C) The next eigenvector *e*_3_, corresponding to *λ*_3_ = 0.1974, brings out the anterior-posterior separation. The anterior and posterior regions are partitioned, respectively, by negative and positive coefficients of *e*_3_. (D) The next eigenvector *e*_4_ corresponds to *λ*_4_ = 0.2836, and it extracts the hemispheric and anterior-posterior partitions simultaneously. Further eigenvectors bring out increasingly smaller structural partitions.

### Fixed points of the connectome

The stable fixed points of the network 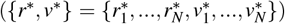, that represent the brain states, are numerically calculated. The details of this calculation are given in the methods section. While each network state is a 2N-dimensional vector, there is redundancy in the sense that, for each region, both variables exhibit qualitatively similar behavior. Hence we limit our quantification of system dynamics to the dynamics of the membrane potential without any loss of generality.

Based on this, the network states are quantified using two scalar metrics, namely *composition* and *stability*. The *composition* of a network state is defined as the number of regions in the up-state and takes integer values between 0 and *N*. The *stability*(*λ*) of a state is defined as the magnitude of the real-part of the largest eigenvalue of the Jacobian of the system, evaluated at the fixed point. A negative value implies that the state is fully stable while a positive value indicates that the state is partially unstable in at least one component. The results show that for significant values of coupling strength, states with higher compositions easily lose stability, Fig. 3(A,B). The *global coupling strength* (*g*), which regulates the cumulative effect of the connectome, quantifies the possible set of dynamical states, which in the symmetric case (*g* = 0) spans 2^*N*^ states. An increased role of the structure constrains the dynamical repertoire of the BNM. The contraction of the sample-space can be explained by understanding the association between *composition* and *stability* of brain states.

**Figure 3.**
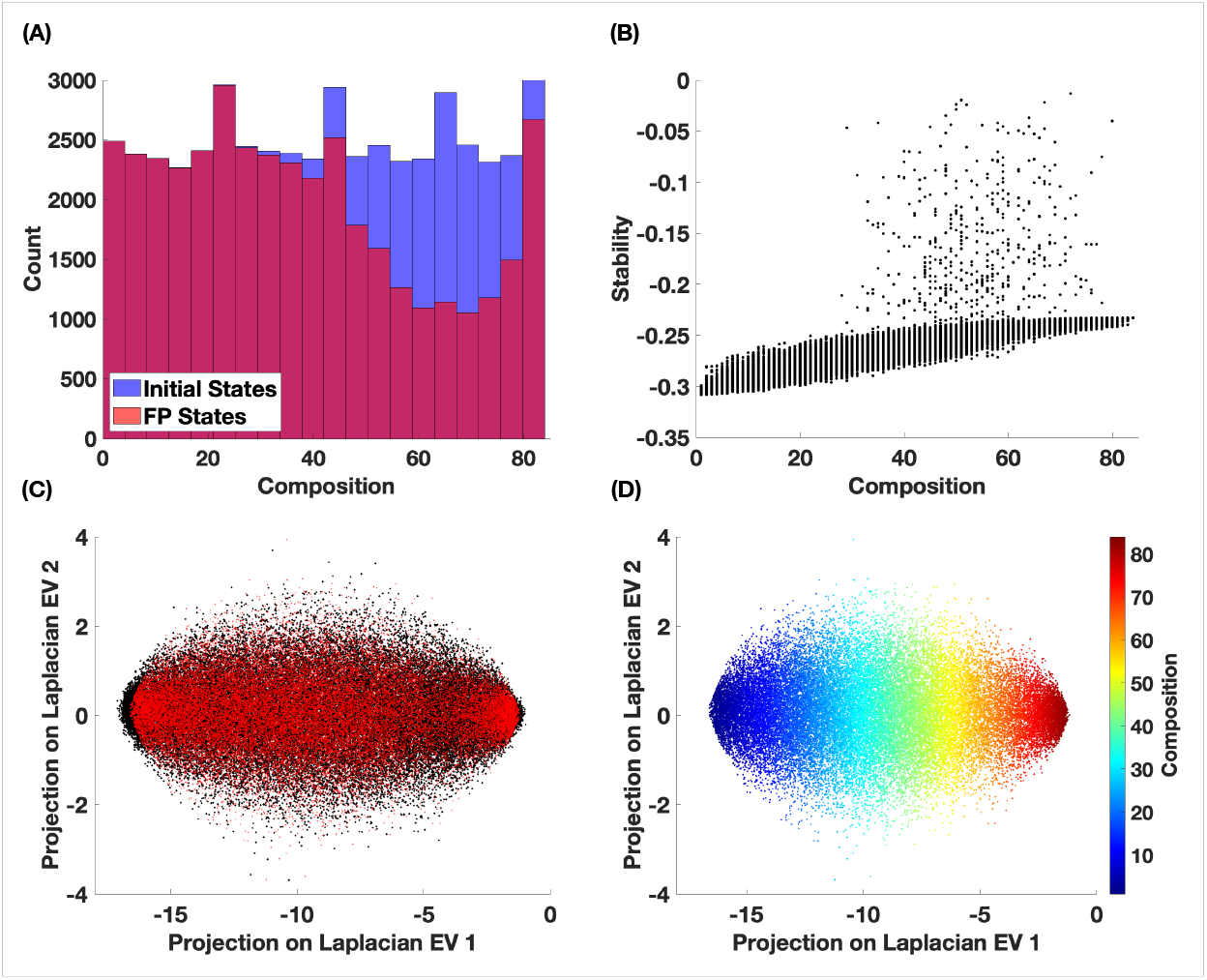
Defining the RS manifold. (A) Comparison of the distributions of compositions of an ensemble of 50000 initial conditions and stable network fixed points for *g* = 0.55. (B) Plot of composition versus stability of the stable fixed points, showing the decrease in stability at higher compositions. (C) Projection of the numerically calculated stable states, relative to the initial conditions, onto the two leading eigenvectors of *L*. The initial conditions and stable fixed point states are colored black and red respectively. (D) Organization of the network states, by composition, on the resting state manifold.

### Structural asymmetry constrains the dynamical repertoire of the BNM

Consider a network state with *composition C*, that has a corresponding *stability λ*_*C*_. When the network is at a fixed point, each region (component) can be treated independently from all other regions, which implies that one unstable region translates to an unstable brain state. The dynamics of the *i*^*th*^ region in a brain state are given by Eqn-9. Assuming that the network has an average in-strength *k*, the interaction term in Eqn-9 can be written as 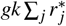. The assumption of an average in-strength holds validity because the in-strengths are quite narrowly distributed around an average value. The fixed-point states of individual regions 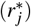 can be coarse-grained into *up* 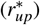 and *down* 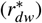 states. Then, for a state with *composition C*, the interaction term becomes 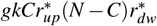. This can be further simplified to *gkC* because the values of 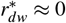 and 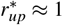, as seen in Fig-5. The interaction term, for each region, can now be treated as an input current term that depends directly on the composition *C* of the global state. Going back to Fig-5, we now have the relationship between the composition of a brain state and the dispersion of individual regions from their respective symmetric-case fixed-point values. More specifically, the deviations grow proportionally with the composition.

Next, if we consider the stability of the *j*^*th*^ region in a brain state, the real-part of the eigenvalue is equal to 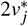. From Fig-5(A), we observe that in the up-state, *v*^*\**^ *≈* 0 but negative; while in the down-state, *v*^*\**^ *≈ −*2. Therefore, between the two, the up-state is significantly less stable than the down-state. And Fig-5(B) indicates that an increasing input pushes 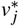 closer towards 0, especially in the up-state. Hence, for any given region, a higher composition of the network state leads to higher regional inputs, thus resulting in lower stability. Between the up and down states, the up-state is more susceptible to this effect.

Therefore, if a global state has a higher composition, then there are more regions in the up-state that receive a higher input, thus lowering the regional stability more quickly compared to a state with a lower composition. This explains the observations for *g >* 0, in Fig-4(B), where the states that quickly lose stability are those with higher compositions. For small values of *g*, the above effect is limited to states with very high compositions. But as *g* is increased, regions in the down-state get pushed to the up-state and the same process continues, and this explains the observations in Fig-4(C).

**Figure 4.**
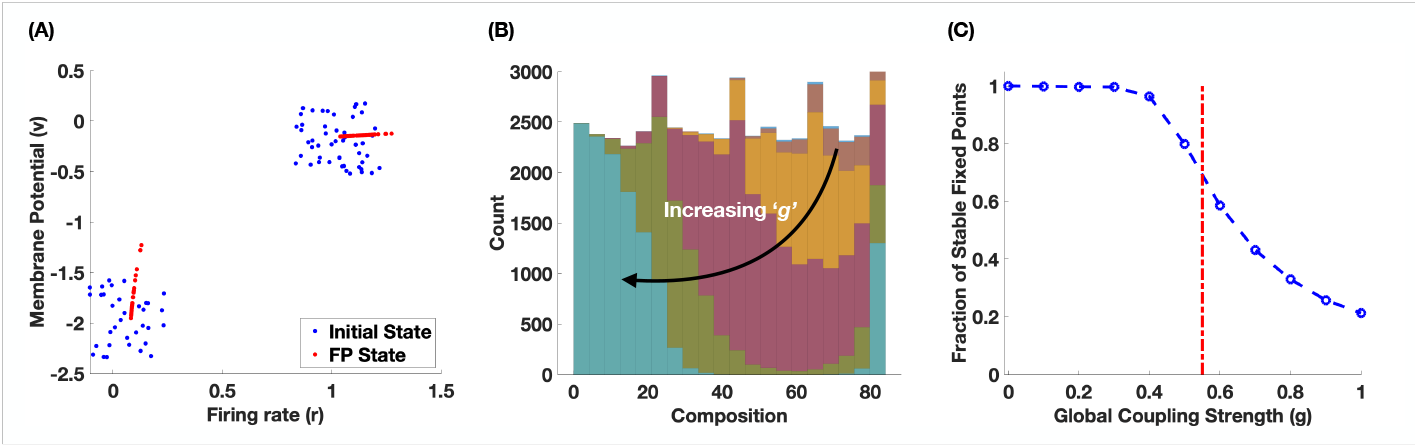
Calculation of fixed-point states. (A) Components of an initial condition with randomly chosen composition and the corresponding fixed point. (B) Change in the distribution of compositions of stable fixed points of the system for decreasing values of global coupling strength. (C) Decrease in the fraction of stable fixed points as a function of global coupling strength.

**Figure 5.**
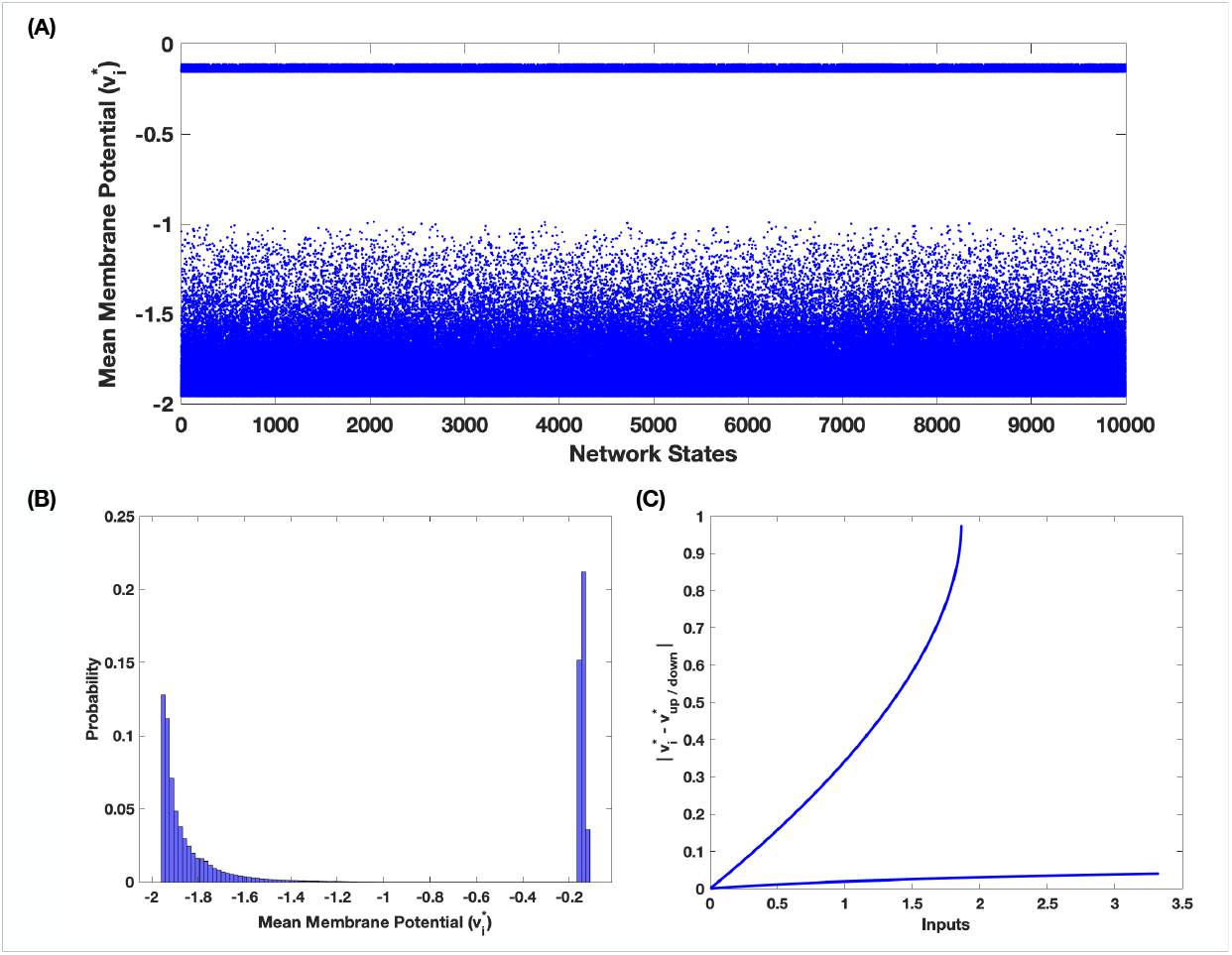
Leading up to Degeneracy. (A) Scatter plot of all numerically calculated network fixed points (only v variable). (B) Distribution of fixed-point components of all network states. (C) Deviation of the fixed-point components from the symmetric-case fixed point as a function of input. The deviations in the up/down state correspond to curves with lower/higher slopes.

### Resting State Manifold (RSM)

A low dimensional representation of brain states, that we refer to as the RSM, is constructed by projecting the fixed points of the system onto the leading eigenvectors of the Laplacian matrix. The details of this process are given Methods-3. This projection nicely separates the network states and spans the manifold. The projection is not unique and others exist^51^, but it is convenient. Based on our knowledge of the eigenvectors of *L* (Fig. 2), we observe along the horizontal axis, a smooth left-right gradient in the composition of states (Fig. 3D) and on the vertical axis, is the separation of hemispheres.

In this new representation, it is easy to visualize how the stable fixed points of the system define a sample space or set of possible configurations for the network. In response to any perturbations (noise or stimulating currents), the system traces a trajectory in this space, as the perturbation induces a switching behavior which in turn is the consequence of symmetry-breaking by the network structure.

### Degeneracy

As mentioned in the previous section, given a system of *N* identical nodes connected by a network *W*, the complex topology introduces a symmetry-breaking in the system. In the symmetric case, where the regions are either isolated or identically connected, all regions in the up-state reach the same fixed point 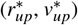 and in the down-state, they all go to 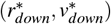. For *g >* 0, this symmetry is broken as each region receives a unique input, that is a function of the topology (in-strength) and the existing configuration of the network. Regions that receive different inputs go to slightly different fixed points and this excursion from the symmetric fixed point is directly proportional to the input. The manifestation of symmetry-breaking and the connection to the framework of degeneracy are depicted in the following figure.

As seen from Fig. 5C, if a region in the down-state receives high-input, the new location of the fixed point moves closer to the up-state and vice-versa. This implies that each region has a different likelihood of transitioning to the other state, depending on the input at that instant. This effect of symmetry-breaking, that results in a differential response to the same perturbation, is transformed and interpreted in the framework of the degeneracy of components of the network state. In the remainder of this section, we will discuss the definition, calculation, and results pertaining to the degeneracy at the regional and global level.

Degeneracy is defined at the regional level and for a specified parameter of the system and is subsequently extended to network states. Consider parameter *η*, for region *i*. The degeneracy of *i* w.r.t *η* is defined as the absolute maximum excursion, relative to its initial value, that is required to switch the region from its local state. If *η*_*init*_ and *η*_*final*_ are the initial and final values of the parameter respectively, then the degeneracy is given by

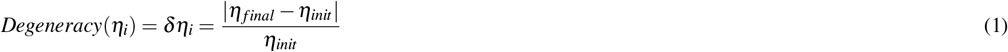

Although the degeneracy is defined as the absolute difference, in certain sections of this work, the sign is taken into account. This is primarily to differentiate the roles of the local states. Further, if a region is in the up-state, the value of the parameter is decreased in order to induce a transition to the down-state and vice versa. This results in upper/lower-bounds of the parameter for regions in the down/up-states. Henceforth, upper/lower-bounds and down/up-states will be used synonymously. The procedure for the calculation of degeneracy is discussed in the methods section.

As a first step in understanding the distributions of degeneracies, only the role of the local state is considered. In Fig. 6 (A - C), the distributions of *δη*_*i*_, *δJ*_*i*_, and *δ*Δ_*i*_ are shown, distinguished only by the local states and independent of regional properties and global states. From Fig. 6(A), it is immediately evident that the distributions for *δη*_*i*_ are distinctly different for up and down states. In the down-state, regions are more likely to have higher values of *δη*_*i*_, whereas they are more likely to have lower degeneracy w.r.t *η* in the up-state. However, in both cases, the range of *δη*_*i*_ is approximately the same. The distributions of *δJ*_*i*_ are qualitatively similar to those of *δη*_*i*_. The actual range of values are, however, well-separated for up and down states. For regions in the down-state, the distribution peaks at much higher values of *δJ*_*i*_. This can be attributed to the scaling by the local state (*Jr*_*i*_ terms) in the network and is discussed in greater detail in the section on geometric interpretation.For degeneracy w.r.t Δ, regions in the down-states are again more likely to have higher degeneracy, while regions in the up-state have no visible distribution and are concentrated in a narrow range on the lower end.

**Figure 6.**
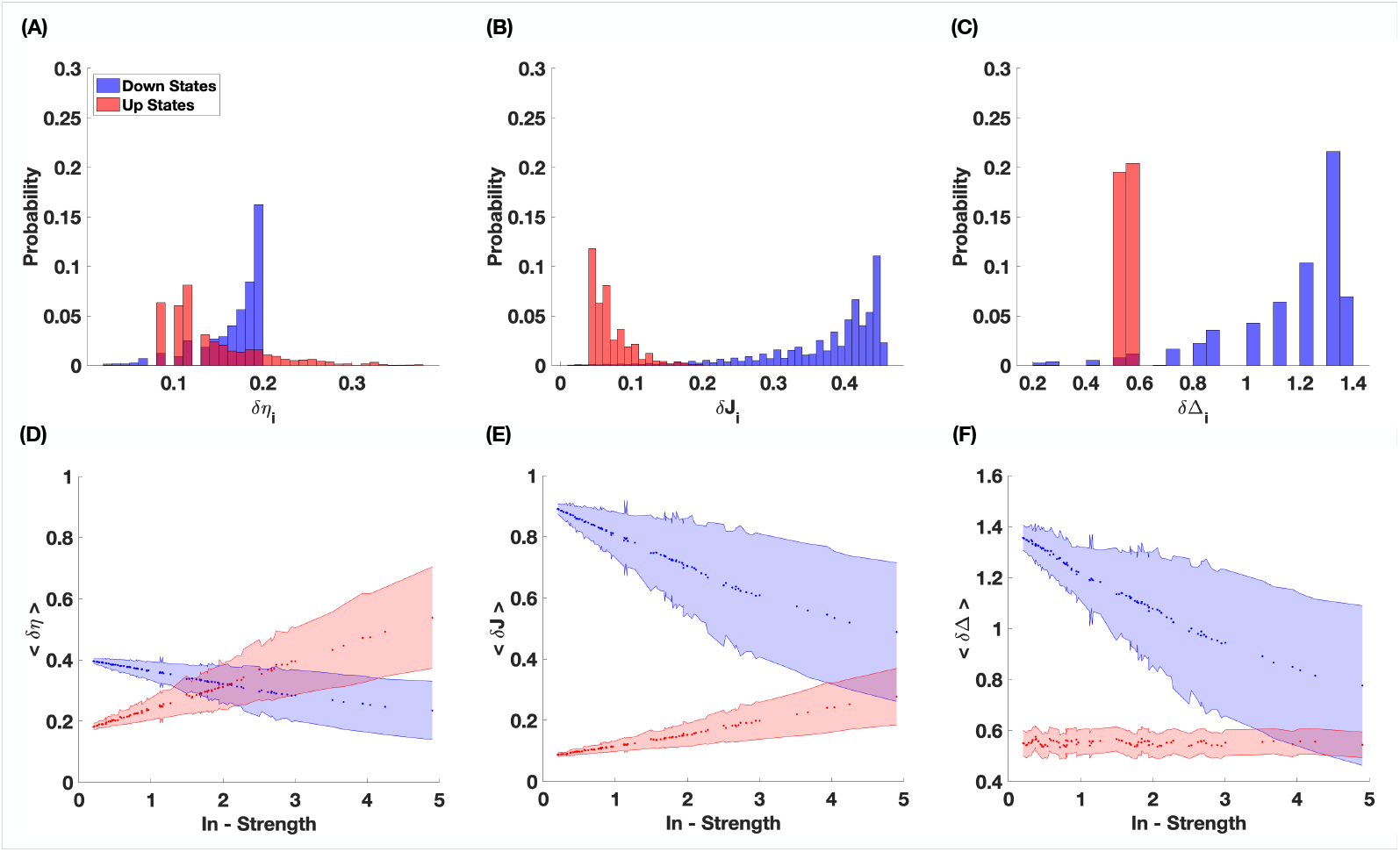
Degeneracy at the regional level. After calculating the degeneracy w.r.t all three parameters, we study the role of local-states and the effect of in-strength. (A - C) Distributions of *δη*_*i*_, *δJ*_*i*_ and *δ*Δ_*i*_, respectively, and the distinction based on local states of the regions. (D - F) The dependence of average degeneracy on in-strength, together with the separation of local states.

Since the degeneracy is a manifestation of the asymmetry introduced by network structure, it is important to understand exactly how it is influenced by the structural properties. The local states, together with the network structure, define the perturbations to the locations of fixed points of individual regions, relative to the symmetric (*g* = 0) case. These deviations significantly affect the response to perturbations, a.k.a degeneracy, and consequently the switching behavior. We focus on the relationship between average degeneracy and the in-strength of the regions. In order to do this, the values of *δη*_*i*_, *δJ*_*i*_ and *δ*Δ_*i*_ are each averaged over all global states for all *i* and plotted as functions of in-strengths of the respective regions and shown in Fig. 6(D - F).

In the down-state, the mean degeneracy decreases linearly with in-strength for all 3 parameters. And for regions in the up-state, it increases linearly with increasing in-strength, except in the case of *< δ*Δ *>*, where it takes a low value and remains independent of in-strength. Therefore, for regions with low connectivity, the local state plays an important role in defining their degeneracy and hence also their response to perturbation. Regions in the up-state are less degenerate and therefore respond more easily to perturbation while in the down-state, they have higher resilience. For regions with intermediate connectivity, the local state plays an important role in the case of *< δJ >* and *< δ*Δ *>*. For *< δη >*, the response to perturbation cannot be predicted based on the local state as the degeneracy takes the same range of values. For regions with high connectivity, there is a greater spread in degeneracy as is observed from the increasing size of variance. Due to this overlapping of degeneracies, it is hard to make the case for a distinct role of local states, for such brain regions. In the next section, we will see how this relationship with in-strength affects the switching behavior as a response to white noise.

So far, we have limited our discussion of degeneracy to the regional level. Each region has been considered in isolation, without concern for the global configuration of the network. Now, we develop two simple approaches to extend the measure of degeneracy to the global state. In both cases, we look for a scalar representation of the network. In this manner, we can study the distributions of degeneracy across the manifold. This allows us to identify subsets of stable and not-so-stable states and their locations, thus laying the foundation for predicting regions with high occupancy and possible transitions between different parts of the manifold in response to perturbations.

In the first approach, the degeneracy (w.r.t each parameter) of the global state is defined as the minimum of absolute values of the degeneracy of all regions (*min*(|*δη*_*i*_|, *min*(|*δJ*_*i*_|, *min*(|*δ*Δ_*i*_|), *∀i*). Once the minimum value has been identified, the local state of the selected region is accounted for by assigning a +*/−* for down/up states (Fig. 7(A-C)). By assigning the minimum value, we identify, for each global state, the most vulnerable component that is most likely to flip its state, hence changing the global state itself. From an alternate perspective, it provides a lower limit for the amplitude of perturbation, that would be required to induce switching.

**Figure 7.**
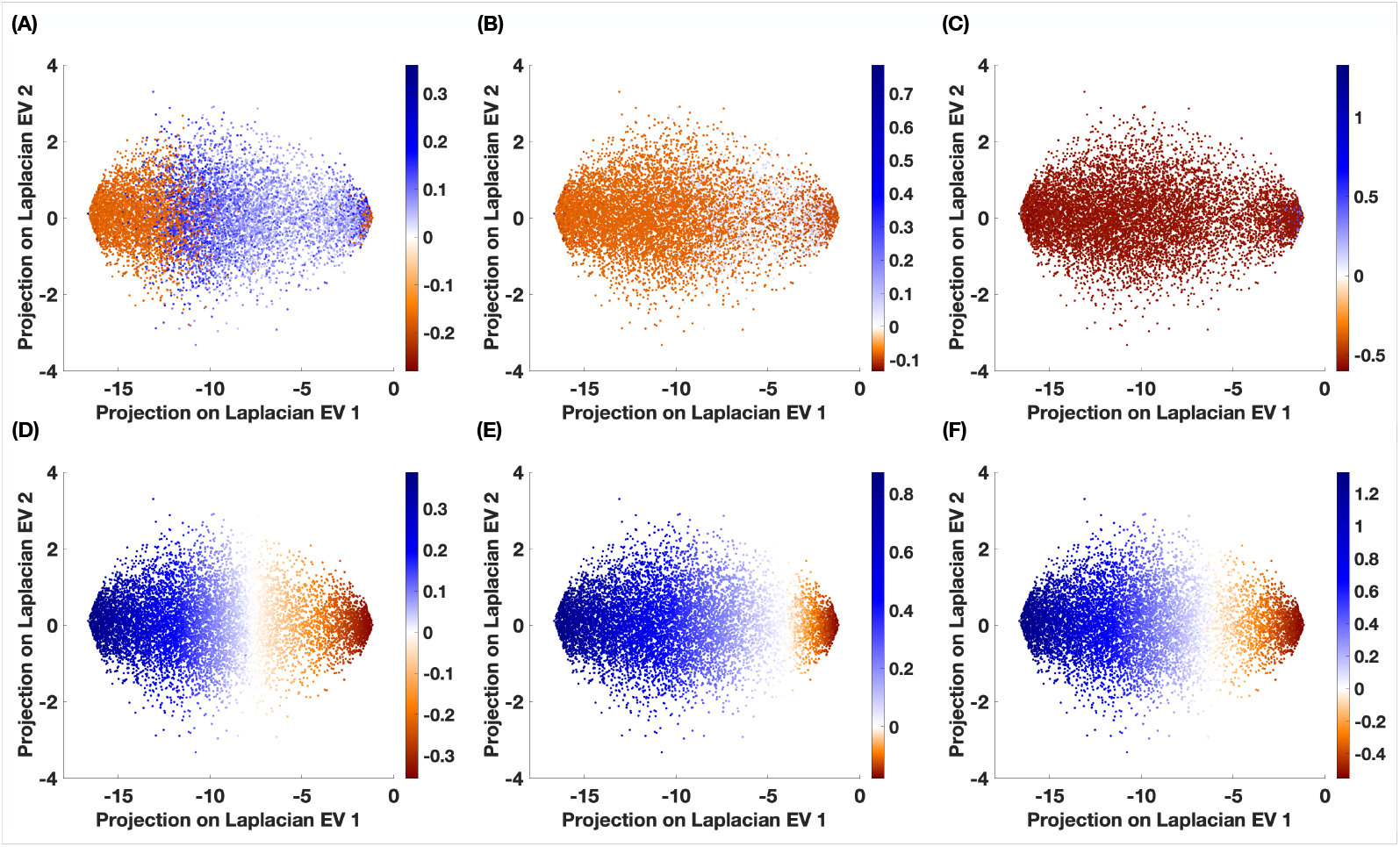
Degeneracy at the global level. Numerically calculated stable fixed-point states are plotted on the resting state manifold and represented by each of the two definitions of global degeneracy. In A - C, each global state is represented by *min*(|*δη*_*i*_|), *min*(|*δJ*_*i*_|) and *min*(|*δ*Δ_*i*_|) respectively. And in D - F, the states are represented by the mean values of degeneracy w.r.t each parameter respectively. In all cases, the values are placed on the blue to red color scheme. Negative values of minimum/mean degeneracy indicate that the main contribution to the global value comes from a region(s) in the up-state and positive values correspond to contributions of regions in the down-state.

Starting with the distribution of *min*(|*δη*_*i*_|), on either ends of the manifold, the least degenerate regions are in the up-state while in the center of the manifold, it is regions in the down-state that have the smallest degeneracy. As we start from the left-end of the manifold and move towards the center, we find that the absolute value of degeneracy increases very slowly until regions in the down-state start to appear. This is marked by the appearance of the red-points. Over here, the regions already have higher degeneracy and this begins to decrease as we move further to the right, at which point the minimum degeneracy reaches zero. And further to the other end of the manifold, the values increase again. In case of *min*(|*δJ*_*i*_|) and *min*(|*δ*Δ_*i*_|), the least degenerate states are all largely associated with regions in the up-state, with very little contribution from the down-states.

A second approach is to assign to the global-state, the mean value calculated over all regional degeneracies. This provides an average measure of the expected response to perturbation and is particularly relevant when there is a stochastic element to the perturbation. As we observe from Fig. 7(D-F), w.r.t all three parameters, there is a very similar pattern of degeneracy. Along the horizontal axis, the mean degeneracy is high on one side and towards the center of the manifold, it goes to zero and then starts to increase again towards the opposite end. However, on the left-side of the manifold, it is positive due to the dominance of low composition states. Similarly, the opposite side of the manifold consists of high composition states and therefore, we have high degeneracy but with negative values. For each parameter, the balance between degeneracies of up/down states occurs on different sections of the manifold.

In both approaches, it is immediately evident that the gradient of degeneracy is primarily along the axis of the leading eigenvector. Looking back on Fig. 3(D), we see that composition also follows the same pattern, indicating a possible relationship between the two. Therefore, in Fig. 8, we associate the minimum/mean degeneracy to the composition and study the connection between the two global properties. The benefit of this approach is that all discussions concerning the response to a perturbation can now be framed in terms of transitions between states of different compositions.

**Figure 8.**
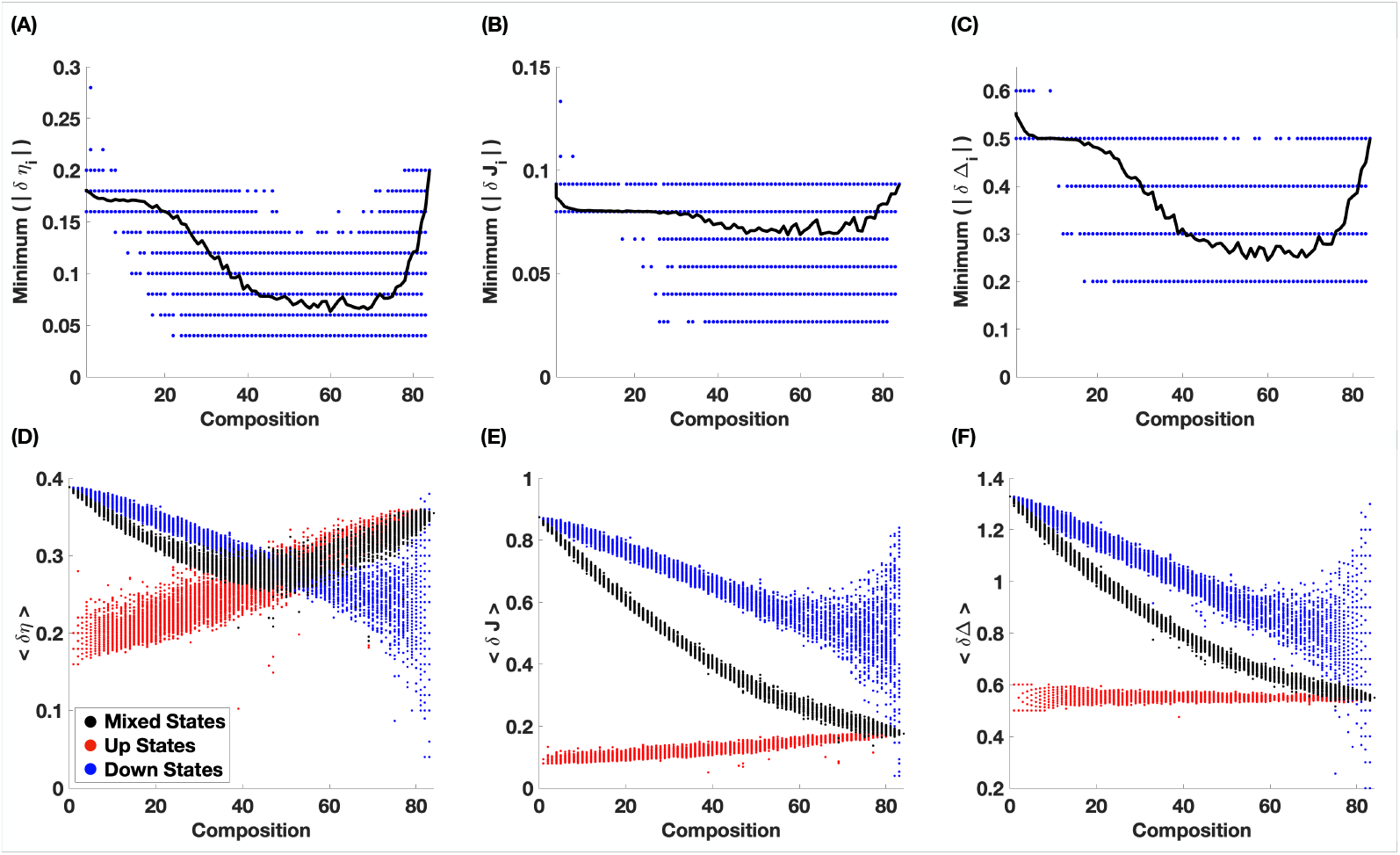
Degeneracy versus Composition at the global level. For each global state, the minimum/mean degeneracy is plotted against the composition of the state. This is shown for *min*(|*δη*_*i*_|), *min*(|*δJ*_*i*_|) and *min*(|*δ*Δ_*i*_ |), respectively from A - C. The unique values of degeneracy for each state are shown in red while the average values of degeneracy of all states with a given composition are shown in black. From, D - F, the mean degeneracy w.r.t each parameter is plotted against the composition of the state. Here, we study three cases, where we consider only the regions in the up-state (red), only regions in the down-state (blue), and the entire global state (black).

From Fig. 7(A - C), we see how the minimum value of degeneracy changes with the composition of the state. On average, for low and high compositions, the degeneracy is relatively high. For states with intermediate compositions, there is a fairly sharp dip (although it is not very pronounced in case of *J*). On the lower end, the average minimum degeneracy is almost constant, pointing to a limited role of perturbations in this range. If regions in a low composition (typically *<* 25) state are perturbed, they switch to the respective opposite states, thus increasing/decreasing the composition on either side. However, the new state also has similar degeneracy, and therefore there is no net preferred direction for the change of composition. At a slightly higher range, the average minimum degeneracy decreases with increasing composition. In this case, if the composition increases, the new state has lower degeneracy and is more susceptible to perturbation. However, if the new state has a lower composition, it is more resilient and therefore, there is a net preference for decreasing the composition. Towards higher compositions, the degeneracy increases rapidly with composition. Following the earlier argument, we should expect a preferential movement towards higher compositions.

So far, we have studied the patterns of degeneracy w.r.t all three parameters of the model. We have looked at the role of local states, the relationship to structural properties and the gradients on the manifold. This distribution of degeneracy defines the contours of stability in the sample-space of fixed-point states of the network. In the following sections, we will narrow down the choice of parameters using a geometric interpretation of degeneracy, with the ultimate goal of understanding and predicting the occupancy and likely transitions on the manifold in a stochastic environment.

### Geometric interpretation of degeneracy

In this section, we briefly steer away from the calculation and consequences of degeneracy and focus on the underlying changes in the phase-space of individual regions. This geometric interpretation allows us to take an analytic approach to degeneracy and opens it up for more general interpretation. Starting from Eqn. 7, the fixed points of the network 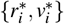 are calculated for a given set of parameters (*η*_0_, *J*_0_, Δ_0_), by solving the following system of equations.

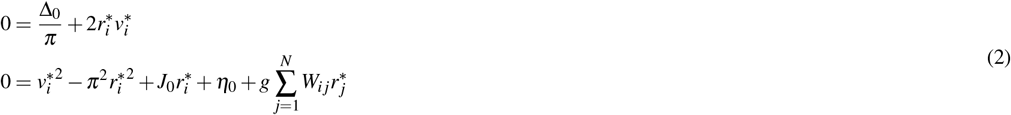

The above system has homogeneous distributions for all 3 parameters and each region goes to a unique fixed point value depending on the respective initial condition and network input. However, the local fixed-point states can be coarse-grained into 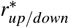 and 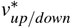. This allows us to categorize each region into up and down states, which then makes it possible to bring degeneracy into the discussion. If we consider parameter *η*_0_ and let *δη*_*i*_ be the degeneracy in *η* for region *i*, then Eqn. 2 can be rewritten to include the degeneracy.

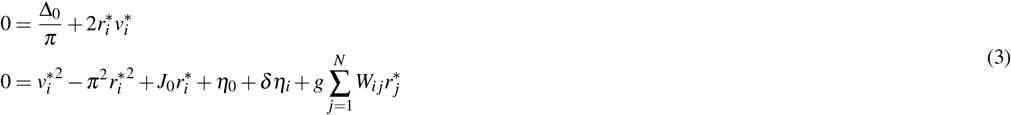

When the network is in a fixed-point state, each region can be treated as an isolated node with a network input term 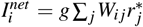. And the entire set of regions can be treated as a set of independent nodes receiving a heterogeneous distribution of input currents 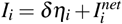. Therefore, for the *i*^*th*^ region, the fixed-point states can be calculated from

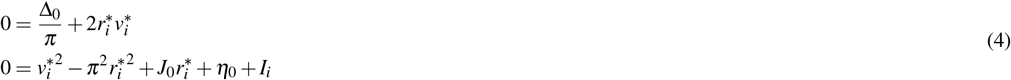

Similarly, if we are interested in *J*_*o*_, and *δJ*_*i*_ is the corresponding degeneracy of the *i*^*th*^ node, then the system of equations can be recast as

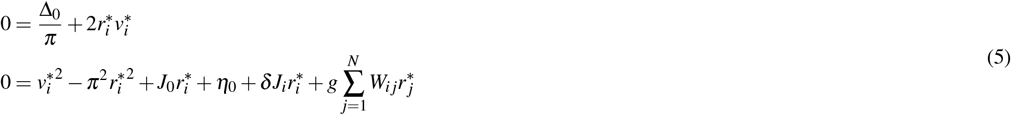

where 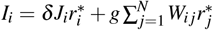. Recasting the equations for degeneracy in Δ results in a more complicated form, but the principle remains just as valid.

Based on Eqn. 7, we can study the effect of the *I*_*i*_ term, as shown in Fig. 9. When *I*_*i*_ = 0, any degeneracy w.r.t a parameter is offset by the input received from the neighborhood. If this is taken as a reference, then the node has a basin of attraction of a particular size, and the up/down states are well-separated. When *I*_*i*_ becomes negative, the size of the basin of attraction of the up-state shrinks rapidly, bringing the stable focus closer to the unstable fixed point. As a result, such regions are constrained to the down-state. On the other hand, when *I*_*i*_ *>* 0, the size of the basin increases, and brings the stable node closer to the unstable fixed point. In this case, the regions are more likely to be locked in the up-state.

**Figure 9.**
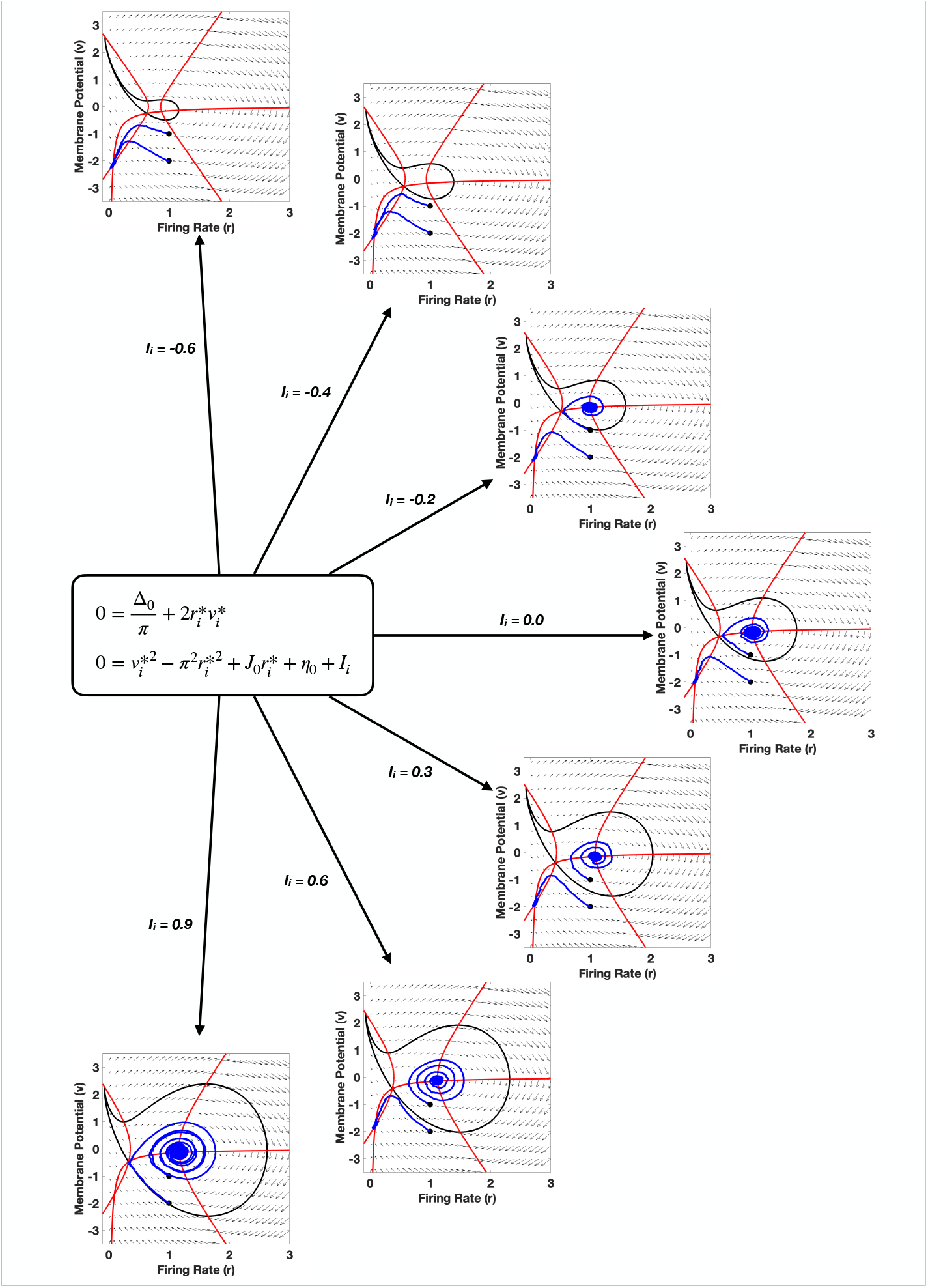
Geometric interpretation of degeneracy. The dynamics of the network in the bistable regime, are plotted in the state space, for different values of input current *I*_*i*_. For each value of *I*_*i*_, we plot the nullclines (red), the basin of attraction (black) and 2 sample trajectories, starting from different initial conditions, to indicate the choice of stable fixed point.

If the symmetry-breaking due to the connectome is to be recast as the degeneracy of the parameters, then *I*_*i*_ = 0 can be treated as the case where the role of the connectome is compensated by the degeneracy of the parameter. Therefore, if a region is in the down-state and has high connectivity, then it is more likely to transition to the up-state than a region with low connectivity. And a region in the up-state, with high in-strength is less likely to transition to the down-state than a region with low in-strength. Alternatively, regions in the down-state with high in-strength have low degeneracy and those with low in-strength have higher degeneracy. And well-connected regions in the up-state have higher degeneracy than those with low connectivity.

Another point worth noting is the relevance of parameter *η*. Comparing Eqns. 6 and 8, we see that the degeneracy w.r.t *η* is not dependent on the local-state, whereas, to relate *δJ*_*i*_ to the in-strength, the local-state has to be distinctly accounted for. For this reason, in the following results, we will depend on *δη* to connect the role of the connectome with the role of degeneracy and study the stochastic exploration of the manifold.

### Noise-driven BNM explores the contours of degeneracy on the RSM

In this section, we study the noise-driven dynamics of the BNM. The description of the simulation is detailed in the methods section. First, we analyze the dynamics at the regional level. Then, we combine this information with our knowledge of the distribution of degeneracy on the manifold, to understand and explain the broader dynamic patterns of the manifold, specifically its occupancy. The low-frequency time-averaged signal is used to study the switching properties at the regional level and the high-resolution source signals are used to study the occupancy and exploration of the manifold. In Fig. 10, for each region, we plot the fraction of time spent in the up-state and the total number of switches versus the in-strength of the nodes. This is done for three cases: low noise, medium noise, and high noise, where the amplitude of noise is controlled by *σ*_*r/v*_.

**Figure 10.**
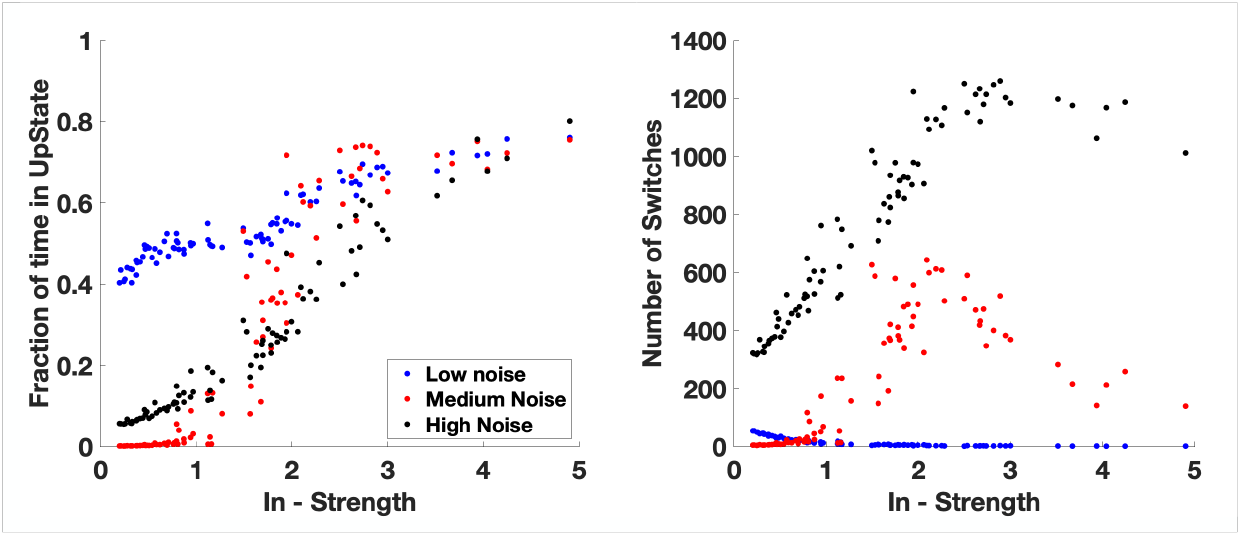
Switching Dynamics. The role of local connectivity on the excitability of regions is studied as a function of in-strength and calculated using the BOLD signal. (A) The fraction of time spent by a region in the up-state is plotted as function of the in-strength of the regions. (B) Plot of number of switches as a function of the in-strength of regions. Both these metrics are plotted for three levels of noise. These two metrics, together, provide the information required to understand the likelihood of individual regions to undergo switching, which in turn lays the groundwork to understand the transition between global states.

As seen from Fig. 10, in the low noise scenario, there is only a quasi-static effect. A majority of the regions remain their respective initial states. Noise-induced switches are few and far in-between and there is no significant time spent in the up-state. This is because the average dispersion due to noise is much lower than the average separation between the up and down states. Hence, low noise has little to no effect on the switching dynamics. Only regions with very low in-strengths show some likelihood for switching. This is further limited to those regions that are initialized in the up-state, which then switch down due to the relatively low stability/degeneracy in the up-state.

In case of high noise, the fraction of time spent in the up-state grows proportionally with in-strength. As for the number of switches, we observe an overall increase. Now, the separation between states is comparable to the amplitude of noise. Hence, switching can occur with relative ease and increases linearly for regions with low in-strength but on the higher end, it remains quite constant. Overall, low noise does not generate much activity and high noise leads almost to random activity in the system.

When the amplitude of the noise is at an intermediate/optimal level, we observe the most interesting behavior. The fraction of time spent in the up-state shows a sigmoidal behavior with regions with low in-strength spending very little time in the up-state while well-connected regions spent the majority of their time in the up-state. This implies that regions with low connectivity are mostly locked in the down-state while those with high connectivity are locked in the up-state. This is consistent with the results from Fig. 6D, where average degeneracy is high for regions in the up-state and with high in-strength and also for those in the down-state and having low connectivity. This lack of activity is further supported by the number of switches, which peaks for regions with intermediate connectivity.

So far, we have looked at how noise plays out at the regional level, given the topology of the network. Using this as a stepping stone, we now look at results on the global level (Fig. 11). A noise-driven exploration of the manifold is studied for three levels of noise and in each case, the system is initialized on different parts of the manifold, corresponding to low, intermediate and high composition states. In each situation, we study how noise impedes/facilitates the ability of the system to explore the manifold.

**Figure 11.**
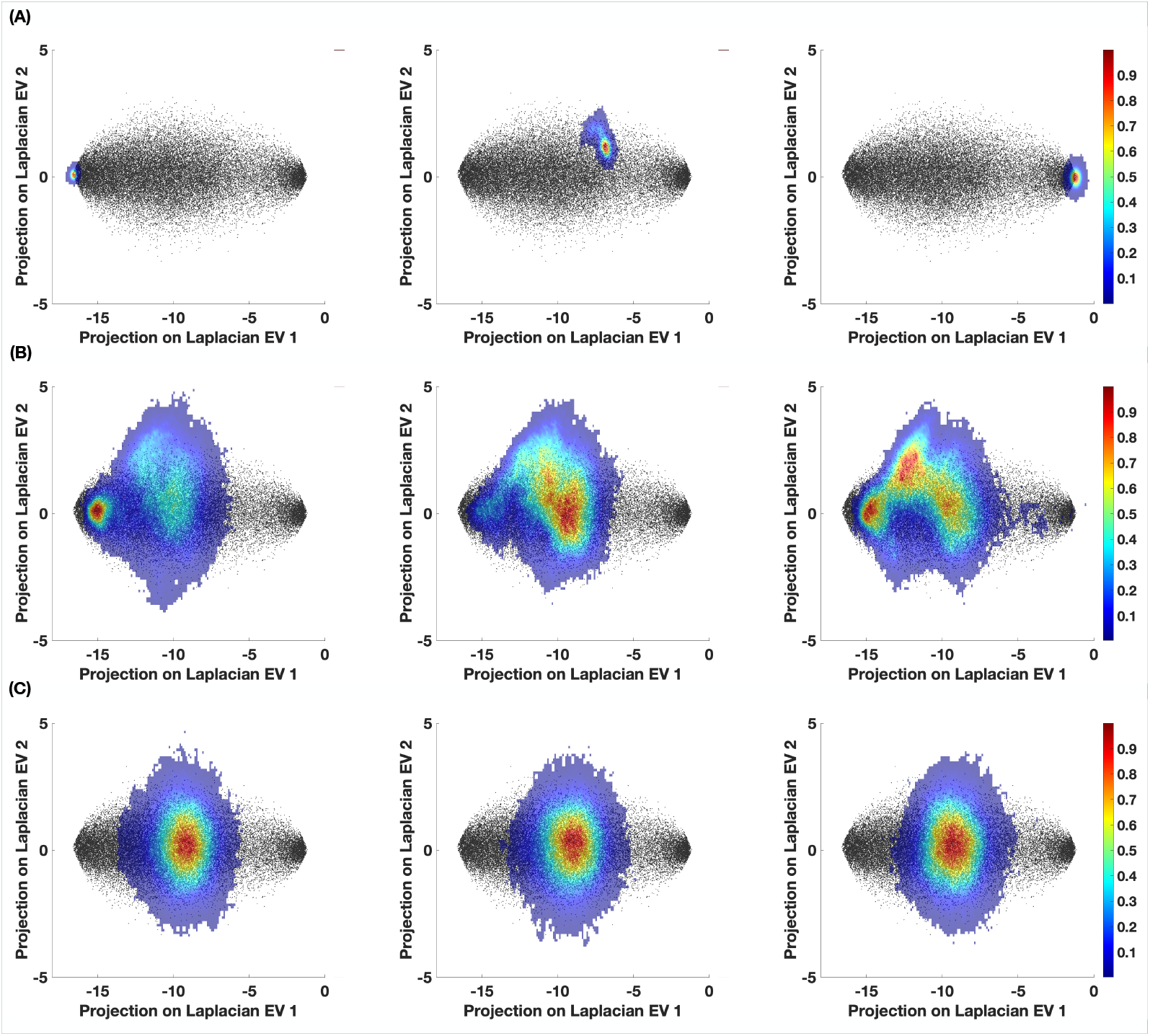
Stochastic exploration of the manifold. The contours of degeneracy on the manifold play an important role in the exploration and occupancy on the manifold. Transitions between various global states are guided by the patterns of degeneracy. The skeleton of stable, analytic fixed-point states, calculated numerically, is plotted in the background in black. The results of stochastic simulation of the system, for (A) low, (B) intermediate and (C) high noise, are plotted as 2D histograms. The color indicates the density of points and is a measure of the occupancy of the manifold. For each category of noise, we consider three different initializations of the network, corresponding to low, intermediate and high compositions (from left to right).

In an environment with low noise, the perturbation is not strong enough to induce switching at the regional level. And the composition of the network remains almost constant, independent of the pattern of degeneracy on the manifold. This is seen in Fig. 11A, where we initialize the system from various locations on the manifold, each corresponding to a different composition. The network does not explore beyond the immediate vicinity of the initial configuration and this is further supported by the distribution of compositions.

On the other extreme, when the amplitude of noise is high, the system always explores the same region of the manifold, irrespective of its initial configuration thus rendering the contours of degeneracy on the manifold largely irrelevant (Fig. 11C). Independent of the compositions at which the system is initialized, it exhibits a symmetric exploration of the manifold. The location on the manifold to which the network converges is where the global mean degeneracy is at a minimum (Fig. 8D). In this part of the manifold, both up and down states have low degeneracy, thus increasing the fraction of vulnerable regions and the overall response to noise. The system shows maximum occupancy in the part of the manifold, that corresponds to compositions spanning half the size of the network, and drops off exponentially for higher/lower compositions. In other words, the exploration is completely random (dominated by noise) and the behavior can be approximated by the average features.

However, an intermediate amount of noise presents the most optimal scenario, in which the exploration of the manifold is maximized. When the network is initialized in a high-composition state, the majority of regions rapidly switch to the down-state due to the less stable nature of the up-state. This leads to an accumulation in the center of the manifold, where global degeneracy is at a minimum. As mentioned earlier, in this part of the manifold, there are a comparable number of vulnerable regions in both states. Therefore, from this point, the composition can either increase or decrease, corresponding to a respective movement to the right or left on the manifold. An increase in composition serves no purpose as the regions quickly switch down again. However, when the composition decreases, the system moves further left, to an area of the manifold where the most vulnerable regions are in the up-state. This drives the system towards even lower compositions. Since the mean degeneracy is higher and the effect of noise is lower, the system spends considerable time in this part of the manifold. Overall, depending on the initialization of the system, it gravitates towards intermediate and low composition states and this is evident from Fig. 11B.

In summary, while there is almost no exploration for low noise, there is limited exploration for high noise. An intermediate amount of noise provides the most optimal setting, which allows the system to occasionally make large jumps between different areas on the manifold, that correspond to transitions between low and intermediate composition states. This correspondence allows us to move beyond the purely spatial features and segue into the temporal aspects of stochastic exploration.

## Discussion

The degeneracy due to the connectome gives another level of complexity and non-identifiability for the structural causes of the brain dynamics at rest^10,34^, in addition to the brain’s multiscale organization, which is fundamental to its dynamic repertoire^56^. In this work, we set out to explore and understand the role of symmetry-breaking due to the structural connectome, and its role in the emergence and exploration of brain states. We show that the asymmetry arising due to the connectome can be captured in the degeneracy of one or more of the system parameters. The patterns of degeneracy, in turn, regulate the dynamics of the broader system in the presence of external and/or internal perturbations. All of the above is realized in the framework of brain network modeling, where the stable fixed points of the network emerge as brain states.

Given a brain network model, the overall effect of the structure quantifies the possible set of dynamical states. Based on a coarse-grained distinction of the firing activity of individual nodes, we defined up and down states. Building further on this distinction of activity on a global level, we introduced the notions of *composition* and *stability*. We also showed that composition and stability are closely linked with the network topology through the properties of individual nodes. While these scalar properties are by no means the most complete descriptors of a brain state, they provide just enough information for us to quantify the effects of structural symmetry breaking.

A low dimensional representation of the brain states, namely the Resting State Manifold, is generated by collapsing the original states onto the two leading eigenvectors of the Laplacian of the structural connectome.^38^ Doing this served three very important goals: first, we can transform very high-dimensional data into a reasonably low dimensional space without significant loss of information at the network level. Second, in the Laplacian space, the brain states are much better organized in terms of our metrics, composition, and stability. Third, and possibly the most important, the eigenvectors of the Laplacian matrix exploit the topology of the connectome to generate a hierarchy of spatial organizations. In the current work, we have only used the 2 leading eigenvectors. However, the hierarchy can be used to extend this approach further. Identifying those eigenvectors that best correlate to well-established sub-networks in the brain, like various lobes, or Resting State Networks, or circuits related to specific tasks. By projecting the original states onto these select eigenvectors, it would be possible to study their specific activation/contribution to the overall activity. This reason also sets apart the choice of a Laplacian eigenbasis from the more commonly used Principal Component Analysis^10,57^.

Defining a symmetric-case fixed-point scenario as a baseline, the strongest effect of the SC appears on the dispersion of individual regions from this scenario. We exploit this effect to cast the topological asymmetry in terms of the degeneracy of the parameters of the BNM. Degeneracy, as defined in Eqn-1, is calculated at the local and global level and its relationship to local connectivity was studied. The contours of degeneracy were mapped on the RSM, which then allowed us to relate it to the composition of the brain states. The observed patterns of degeneracy were suggestive regarding the role of perturbations. In particular, they predict the occupancy of different brain states and consequently the exploration of the manifold. This was validated by using standard Gaussian noise as a perturbation to study the exploration of the manifold and the emergent structured flow on it, for different levels of noise and initializations.

While the framework of degeneracy lends itself to be a valuable tool to understand the layout of the RSM and predict the effect of perturbations (in particular standard normal noise), thus also pointing to brain resilience, an important limitation lies in the coarse-graining of the activity into up and down states. Even though these stem from the basic feature of the neuronal populations – their activity is broadly divided into low- and high-firing rates^58–60^, during the awake state the high-firing rate activity is prominent with shorter periods of lower firing rate (e.g.^61^). However, this same approximation also allows us to extend the understanding of the effects of structural asymmetry in terms of functional/dynamical asymmetry. Essentially, we can draw an equivalence between the role of the structural connectome and the regional heterogeneity in the BNM^62^. Although the latter is known to improve the fit of BNMs^6,33,63^ we have omitted it in our work to focus on the impact of the connectome heterogeneity, which is a more generic feature and better established feature of virtual brains^13^. Further, the results that are obtained by a noise-driven exploration of the RSM, can be directly related to the escape-time analysis described in^10^. Taking it a step further, the distribution of degeneracy, calculated for an optimal parameter combination, can be used to quantify the occupation probability (in place of escape-times) of either individual brain states or states of identical composition. This would open the possibility of understanding the exploration of the manifold in terms of transition probabilities between different states. The results in the case of optimal noise have a significant bearing on the notion of a working point of the BNM. The *working point*, as discussed in^55^, corresponds to the maximization in the exploration of the manifold, or the point of maximal fluidity^10^, and results in a strong correlation between the structural connectivity and the functional connectivity of the simulated activity. Moreover, the same working point of maximal dynamical fluidity has been associated with homeostatic brain activity^18^ that changes during different levels of consciousness^22^, and is closely linked to the concept of high metastability as a desired working point of the brain^20,64–66^.

In conclusion, we have explored the role of symmetry-breaking due to the structural connectome, in a Brain Network Model. It defines the contours on a low dimensional resting state manifold, which in turn guides the stochastic exploration of the space, giving rise to a structured flow on the manifold. This phenomenon is explained in the framework of degeneracy of network states, w.r.t the local parameters. This framework can also be extended to understand the effects of symmetry-breaking in the context of regional variance or spatial heterogeneity.

## Methods

### Construction of the BNM

Dynamics at the regional level are simulated using the neural mass model proposed by Montbrio et al.^35^ in 2015. It is a 2D nonlinear model, derived from an ensemble of all-to-all coupled Quadratic Integrate and Fire (QIF) neurons, and describes their mean-field behavior in a mechanistic manner.

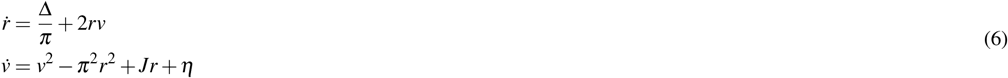

The model, given by Eqn 6, describes the dynamics of the mean firing rate (*r*) and mean membrane potential (*v*). The average connectivity in the ensemble is given by synaptic weight *J*, while the heterogeneity of input currents in this ensemble is modeled by a Lorentzian distribution with mean *η* and half-width Δ. In the phase-plane (Fig. 1A), the system has two stable fixed points: a stable focus and a stable node, that define our *up* and *down* states respectively. In Fig. 1A, we plot the nullclines (in red), the basin of attraction (in black), and the phase-flows (as arrows). Sample trajectories of the system, in the vicinity of both the stable focus and node, are also shown in blue.

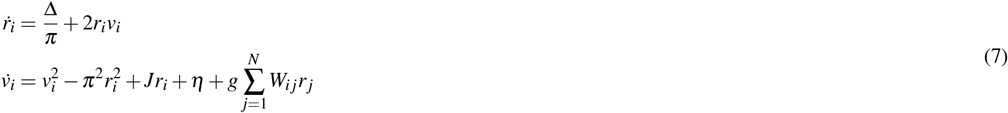

The brain network model (BNM) is constructed, using the HCP-001 structural connectome (*W*) of size *N*, where each brain region is modeled by Eqn. 6. While this is an arbitrary choice of connectome, there is no loss of generality in the results. This connectome is based on the DK atlas, with 84 regions, including both cortical and sub-cortical areas. The values are normalized to [0, 1], which also allows for comparison of results across connectomes of different subjects. All regions are set in the bistable regime with constant parameter values given by Δ = 1, *J* = 15, *η* = *−*5 and are linearly coupled via the firing-rate variable. The effect of network structure is controlled using global coupling strength (*g* = 0.55).

### Laplacian Eigenbasis

While the normalized connectome is used for the calculation of the fixed points and for stochastic simulation, it is the Laplacian matrix that is more interesting in terms of dimensionality reduction. The symmetric normalized form of the Laplacian(*L*), of *W*, is defined and calculated as

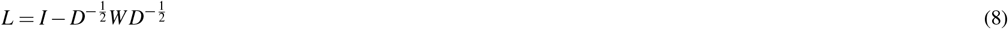

where *D* is a diagonal matrix whose entries are the in-strengths of *W* and *I* is the Identity matrix of order *N*.

#### Calculation of network FPs

The fixed points of the network are numerically calculated using the multidimensional Newton-Raphson algorithm to solve the following equations.

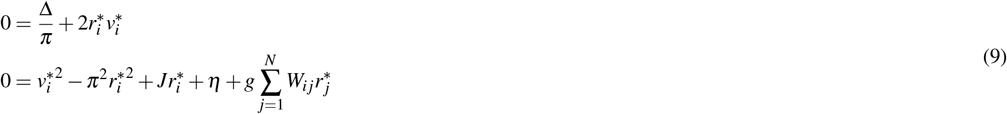

The compositions of the initial conditions are sampled from a uniform distribution and the values are chosen randomly from a small region around the symmetric (uncoupled) case fixed points. The corresponding fixed point and its stability are calculated. The fixed point is retained if it is stable and rejected otherwise. An example is given in Fig. 4(A).

This process is repeated for a set of 50000 initial conditions, for a given value of global coupling strength. For positive values of *g*, the number of stable fixed points starts to decrease. An example of the distributions of initial conditions and the corresponding stable states is shown in Fig. 3(A). For the symmetric case (*g* = 0), it is equivalent to having *N* independent systems and therefore, we can have 2^*N*^ stable states. As *g* becomes more positive, the states with high compositions start to become less stable, and *λ* starts to move closer to zero. This indicates the contribution of high-composition states to the overall decrease in stability. Changes in the distribution of compositions of stable states for increasing *g* are shown in Fig. 4(B). Beyond a certain threshold value of *g*, the fraction of stable fixed points rapidly decreases, as shown in (Fig. 4(C)). This threshold marks the start of our working-point regime, where the system starts to show an increasingly strong correlation between functional and structural connectivity. The FCD also begins to become non-trivial and this is quantified by calculating the variance of the upper-triangular part of the FCD matrix. Based on this understanding of the role of global coupling strength, we set *g* = 0.55 as our working point in this study.

#### Construction of the RSM

Given a network with *N* regions, where each region can exist in either up/down state, it is possible to have 2^*N*^ stable global states in the symmetric (*g* = 0) case. While the number is substantially lower for higher values of *g*, it is still very high-dimensional. This makes it more difficult to visualize, understand and interpret the results, even when the states are represented by a scalar value. Therefore, we need a new representation that reduces the dimensionality of the results while simultaneously improving the visualization and interpretability.

To this end, the eigenbasis of the Laplacian matrix (*L*) lends itself as an ideal candidate. A low-dimensional representation is constructed by projecting the stable fixed points onto the two leading eigenvectors of *L*. This generates a manifold, where each network state is represented by a single point in 2D space (Fig. 3C). In this new space, the states are neatly organized, so that some of the information is also visually evident. Since we are concerned with the properties of the resting state of the human brain, we define it as the resting state manifold.

#### Calculation of degeneracy

The following procedure is used to numerically calculate the value of degeneracy w.r.t a specific parameter. A fixed-point state of the network is selected and the value of the parameter is incrementally changed for each region. After every increment, the system is integrated in a deterministic manner to find the new fixed point of the network. This is followed by a coarse-graining step to bunch the regions into up/down states. Depending on the initial state of the region, the parameter value is either increased or decreased, until it undergoes a transition to the opposite state. It is worth noting that, in a given network state, the region for which the parameter is modified, is the one that is most likely to switch while the other (connected) regions show negligible deviation from their respective fixed points. This process is repeated independently for all regions in all network states, giving us a collection of degeneracy values, that are a function of the local as well as the global states. A demonstration of this procedure, with examples for an up-switch and down-switch, is provided in Fig. 12.

**Figure 12.**
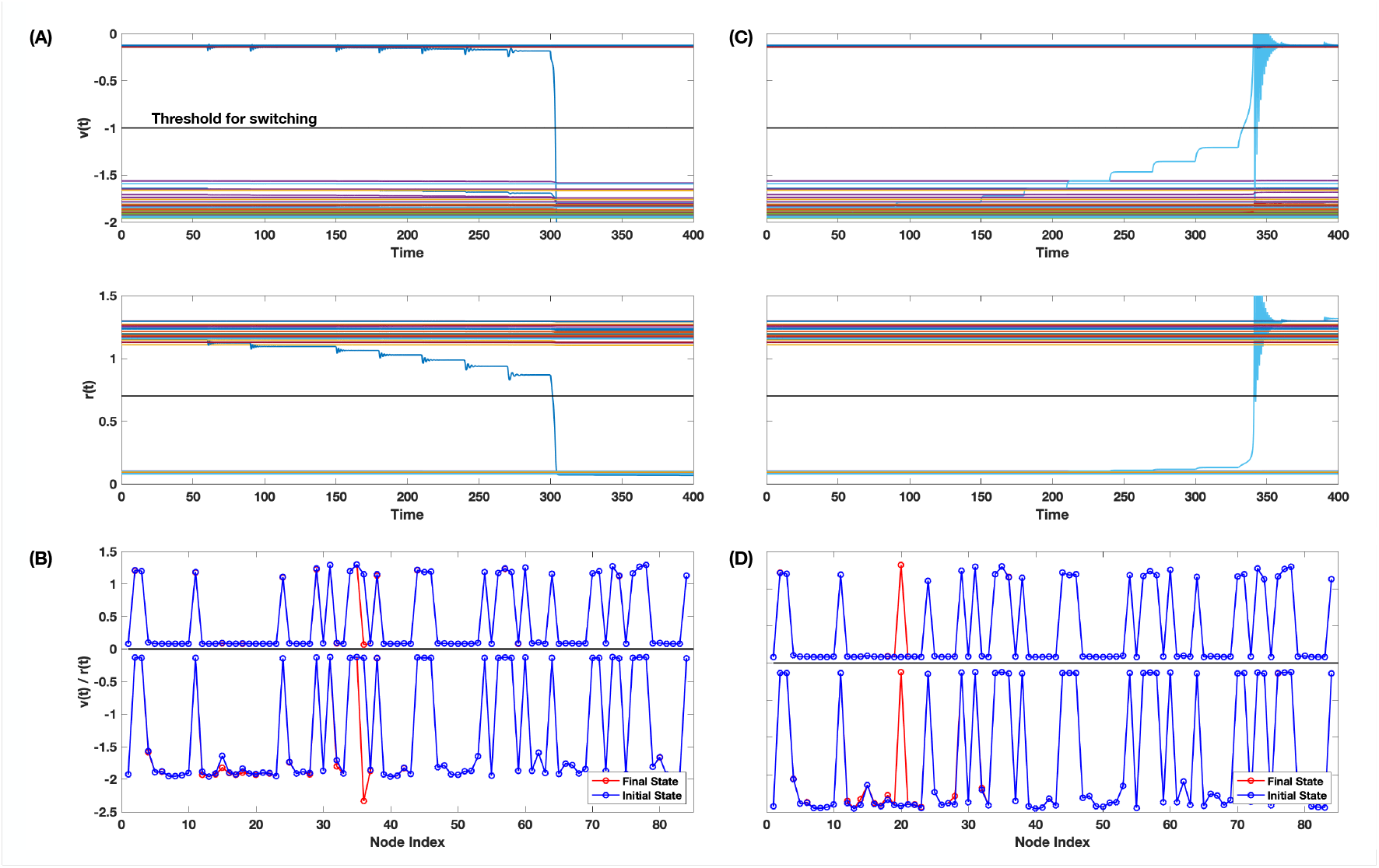
Calculation of Degeneracy. The procedure for calculation of degeneracy is demonstrated using two examples. An initial fixed-point state of the network is arbitrarily selected. Two regions, one each in the up and down states, are chosen. If the selected region is in the up/down-state, the value of *η* is decreased/increased in steps of 0.2, followed by deterministic integration to the new fixed point. Time-series of both variables, corresponding to this sequence of parameter increments and subsequent integrations are shown in (A) and (C). Corresponding initial and final fixed-point states of the network are shown in and (D), clearly indicating the switching of the selected regions, in both variables.

#### Stochastic integration of the BNM

A stochastic simulation of the BNM is performed by integrating the following system of equations using the stochastic Heun method.

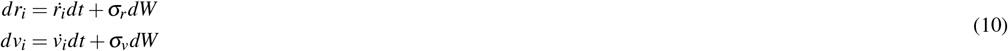

where 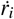 and 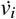 are taken from Eqn - 7, *dW* are independent random variables drawn from the standard normal distribution and *σ*_*r*,*v*_ are the diffusion coefficients scaled respectively by the dynamical ranges of the variables. The values of *σ*_*r*,*v*_ are adjusted to simulate low (*σ*_*v*_ = 2.*σ*_*r*_ = 0.014), medium (*σ*_*v*_ = 2.*σ*_*r*_ = 0.037) and high noise (*σ*_*v*_ = 2.*σ*_*r*_ = 0.051) environments.

Other parameters of the integration include *J* = 15, *η* = *−*5, Δ = 1 and *dt* = 0.01*ms*. In each case, the source signals are generated in 20-minute segments with 1*ms* resolution and a time-averaged low-frequency signal is generated with 2*s* resolution.

